# Homonuclear Chemical Shift Correlation in Solids Under MAS by Fast Cross-Relaxation Driven Spin Diffusion

**DOI:** 10.64898/2025.12.12.694067

**Authors:** Riqiang Fu, Ayyalusamy Ramamoorthy

**Affiliations:** United Imaging NMRSpec Scientific Instrument Co. Ltd, Wuhan, Hubei 430206, China; National High Magnetic Field Laboratory, Florida State University, 1800 East Paul Dirac Drive, Tallahassee, FL32310, USA; Department of Chemical and Biomedical Engineering, FAMU-FSU College of Engineering, Florida State University, 2525 Pottsdamer St., Tallahassee, FL32310, USA; Institute of Molecular Biophysics, Florida State University, 91 Chieftan Way, Tallahassee, FL32304, USA

**Author notes:** Corresponding author: Dr. Riqiang Fu.

**Keywords:** homonuclear chemical shift correlation, Cross relaxation driven spin diffusion, Hartmann-Hahn matching condition under MAS, ^15^N-^15^N correlation

## Abstract

In this study, we report a two-dimensional NMR technique that correlates the chemical shifts of homonuclear spin systems in solids under MAS. The pulse sequence employs double spin-lock RF pulses to facilitate magnetization exchange among low-*γ* nuclei (such as ^13^C or ^15^N) through cross-relaxation driven by a combination of spin diffusion and RF field. We systematically investigate how the efficiency of the magnetization exchange depends on the Hartmann-Hahn mismatch and the MAS frequency. Experimental results obtained from ^13^C-labeled Fmoc-Leucine powder sample, ^15^N-labeled L-histidine amino acid (pH 6.3) powder sample, and uniformly-^15^N-labeled aquaporin reconstituted in DMPC lipid vesicles are reported. The results reveal a rapid spin-exchange process, with transfer rates that qualitatively correlate with the internuclear distances of the participating low-*γ* nuclei. It is remarkable that a 50 ms DARR mixing resulted in no cross peaks in the 2D ^15^N-^15^N chemical shift correlation spectrum of uniformly-^15^N-labeled aquaporin, whereas as short as 5.0 ms mixing duration using the double spin-lock mixing yielded 2D spectrum exhibiting cross peaks between neighboring amino acid residues. Our results demonstrate that the proposed approach can be utilized to enable magnetization exchange between nearby ^15^N or ^13^C nuclei, which is highly desirable for accomplishing resonance assignments in the structural studies of proteins.

## Introduction

Multidimensional NMR experiments that exploit homonuclear chemical-shift correlations constitute an essential component of resonance-assignment strategies for a broad range of molecular systems in both solution and solid states [1-2]. In the solid state, under either static or magic-angle spinning (MAS) conditions, such experiments are commonly implemented by correlating the chemical shifts of low-γ nuclei, most frequently ^13^C and ^15^N. These correlations play a particularly important role in high-resolution structural studies, where they provide the basis for determining intra- and inter-residue connectivities in uniformly ^13^C or ^15^N labeled proteins and peptides [3-7]. However, accomplishing such experiments for ^15^N–^15^N correlations in proteins remains challenging because the dipolar couplings between ^15^N nuclei are exceedingly weak, making direct magnetization exchange inefficient on experimentally practical timescales [8-19].

A significant advance in addressing this limitation was reported by Xu et al., who demonstrated that a spin-lock radio-frequency (RF) field applied to either ^13^C or ^15^N can induce magnetization exchange through cross-relaxation–driven spin diffusion under both static and MAS conditions [20]. The application of an RF field during the mixing period effectively enhances spin diffusion by enabling transitions that are otherwise strongly suppressed in the presence of MAS and small dipolar couplings. Building on this concept, later work showed that applying mismatched Hartmann–Hahn spin-lock RF pulses simultaneously on the proton and ^15^N (or ^13^C) channels can further promote efficient magnetization transfer in oriented solids [21] and in powders under static or MAS conditions [22]. Similarly, it was found by Griffin and coworkers that proton irradiation not only assists the polarization transfer between low-γ nuclei [23] but also enhances the homonuclear polarization transfer rate among low-γ nuclei, the so-called proton assisted recoupling (PAR). [24-27]. These developments collectively emphasized the potential of RF-assisted mixing schemes to overcome the limitations posed by weak homonuclear dipolar couplings.

In the present study, we extend these efforts by investigating a double spin-lock mixing sequence designed to enhance homonuclear magnetization exchange among low-γ nuclei under MAS, as in the PAR experiments. However, although the spin-lock fields used in the PAR experiments were primarily guided by numerical simulations, the optimal experimental conditions were highly dependent upon samples according to the experimentally calibrated conditions.[26] Here, we used a uniformly ^13^C labeled Fmoc-leucine powder sample to systematically characterize the dependence of the magnetization transfer efficiency among ^13^C nuclei as a function of the Hartmann–Hahn matching condition under MAS, providing insight into the mechanisms by which the spin-lock fields modulate the effective Hamiltonian during the mixing period. Our experimental results suggest that the most efficient transfer takes place when *ν*_1*C*_ − *ν*_1*H*_ = ±ν_*r*_, where *ν*_1*C*_ and *ν*_1*H*_ are the spin-lock amplitudes for the ^13^C and ^1^H channels, respectively, and *ν*_*r*_ is the sample spinning rate. Our measurements reveal that the double spin-lock scheme under this condition produces a remarkably rapid spin-exchange process, with transfer rates that qualitatively correlate with the internuclear separations among ^13^C spins. These observations suggest that the method is sensitive to spatial proximity and therefore holds promise for structural studies where distance-dependent transfer is advantageous. Directly applying this double spin-lock condition to the uniformly ^15^N-labeled Aquaporin Z protein sample reconstituted in DMPC lipid vesicles, we obtained 1 2D ^15^N-^15^N chemical shift correlation spectrum with a mixing time as short as 5.0 ms. The appearance of cross peaks between ^15^N nuclei present in the adjacent amino acid residues suggest that this technique will be useful to assign resonances from a uniformly labeled protein sample.

## Materials and Experimental

^13^C-labeled Fmoc-L-leucine and uniformly-^13^C, ^15^N-labeled L-Histidine·HCl·H_2_O, pH 3.3 powder sample were purchased from Cambridge Isotope Laboratories, Inc. (Boston, MA) and used without further purification. The crystalline histidine was dissolved in deionized water and titrated with 0.2 M NaOH to reach pH 6.3, then lyophilized in the vacuum pump. The lyophilized histidine powder was packed into a 3.2 mm MAS rotor for solid-state NMR measurements. Aquaporin Z (AqpZ), a ∼24-kDa integral membrane protein that facilitates water movement across *Escherichia coli* cells with a high rate, was expressed and purified detailed in the literature [28] to obtain a uniformly-^13^C,^15^N-labeled AqpZ sample.

All NMR experiments were carried out on a Bruker NEO 800 MHz NMR console operating at the resonance frequencies of 799.79 MHz for ^1^H, 201.13 MHz for ^13^C, and 81.05 MHz for ^15^N nuclei. A home-built (at NHMFL) 3.2 mm low-E double-resonance biosolids MAS probe [29] was used to carry out the reported solid-state NMR results. The sample spinning rate was controlled by a Bruker pneumatic MAS III unit at 16.0 kHz ± 3 Hz. The radio-frequency (RF) pulse sequences used in this study are shown in Figure 1. As shown in Fig. 1a, proton to ^13^C ramped cross-polarization (CP) with a contact time of 1 ms was used to enhance the sensitivity of ^13^C detection. During CP, a 50 kHz ^1^H spin-lock and a 38 to 56 kHz ramped ^13^C spin-lock were employed for an efficient CP. After CP, the carbonyl carbon magnetization was selectively prepared by using a selective spin-echo sequence (*τ*_d_-180_x_°-*τ*_d_) with the echo delay *τ*_d_ = 62.5 μs and a 180° Gaussian shaped pulse for a duration of 5-rotor periods (i.e. 312.5 μs) applied at the carbonyl carbon resonance frequency. Then, the transverse ^13^CO magnetization was allowed to evolve under the double spin-lock fields simultaneously applied to both ^1^H and ^13^C nuclei. A SPINAL64 decoupling sequence [30] with a ^1^H B_1_ filed of 78.0 kHz was applied during the selective spin-echo sequence (*τ*_d_-180_x_°-*τ*_d_) and during ^13^C signal acquisition. 3072 data points were acquired with a 5 μs dwell time, and 16 scans were used to accumulate the signals with a recycle delay of 3 s. The ^13^C chemical shifts were referenced to the carbonyl carbon resonance of a glycine powder sample at 178.4 ppm, relative to 4,4-dimethyl-4-silapentanesulfonate sodium (DSS) at 0 ppm.

**Fig. 1.**
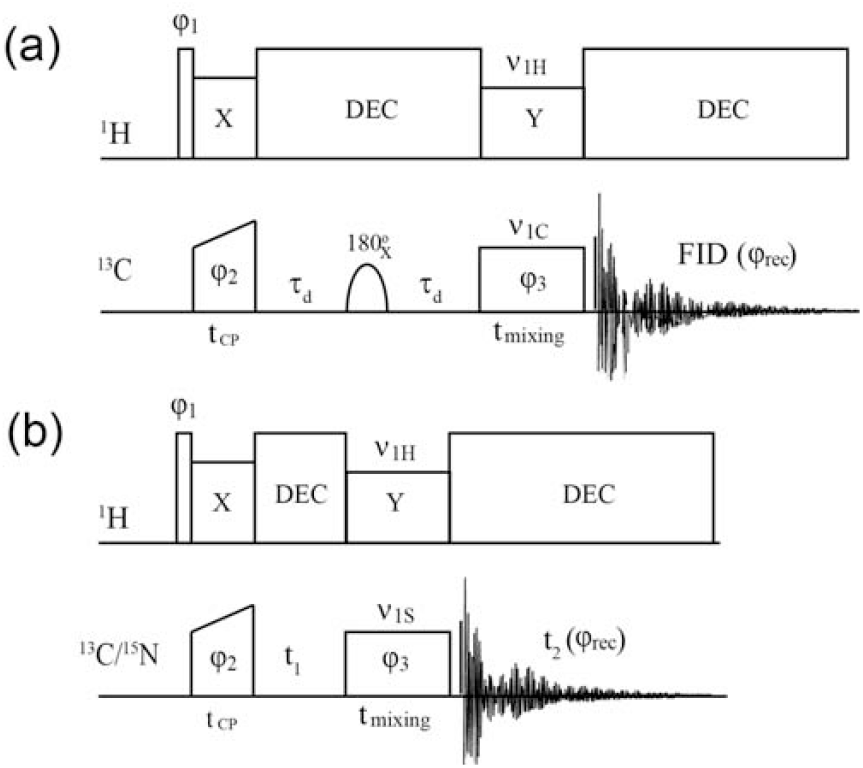
RF pulse sequences used to collect spectra presented in this paper. (a) The 1D scheme to select the carbonyl carbon magnetization and then to monitor its transfer to other ^13^C nuclei by scanning the spin-lock amplitudes *ν*_1H_ and *ν*_1C_. The selective 180° pulse is applied at the carbonyl carbon resonance frequency. The phase cyclings used are: φ_1_ = y -y; φ_2_ = x x y y -x -x -y -y; φ_3_ = x x y y; and φ_rec_ = x -x -y y -x x y -y. (b) A 2D pulse sequence used to acquire homonuclear correlation spectra. The phase cycling used is the same as in (a) except the receiver phase cycling φ_rec_ = x -x y - y-x x -y y.

Fig. 1b shows the pulse sequence used to collect two-dimensional (2D) ^15^N-^15^N chemical shift correlation spectra. In this sequence, the ^15^N magnetization was enhanced by CP with a 2 ms contact time, during which a 50 kHz ^1^H spin-lock and a 38 to 56 kHz ramped ^15^N spin-lock were employed. The spectral widths for the t_2_ (direct) and t_1_ (indirect) dimensions were 50 and 20 kHz, respectively. In the t_2_ dimension, 2048 points were collected. An RF field strength of 62.5 kHz was used in ^15^N RF channel for the dipolar assisted rotational resonance (DARR) experiments, where a ^1^H RF field strength of 16 kHz was used during a 50 ms DARR mixing period. The ^15^N chemical shift was referenced to 33.5 ppm of ammonium nitrogen of a powder sample of glycine, relative to liquid ammonia at 0 ppm.

## Results and Discussion

Fig. 2a shows the ^13^C CPMAS NMR spectrum of a powder sample of Fmoc-L-leucine with the RF carrier frequency set at 109 ppm. The isotropic chemical shifts for the six labeled ^13^C nuclei (i.e. C’, C_*α*_, C_β_, C_*γ*_, C_δ1_, and C_δ2_) were assigned: δ_C’_ = 176.0, δ_C*α*_ = 54.4, δ_Cβ_ = 49.7, δ_C*γ*_ = 27.5, δ_Cδ1_ = 27.1, and δ_Cδ2_ = 23.5 ppm. When the selective 180° pulse was applied to the C’ nuclei, the ^13^C’ signal was refocused, while the other ^13^C signals were completely canceled out after the complete phase cycling. As shown in Fig. 2b, when no spin-locks were applied (i.e., *ν*_1H_ = 0 and *ν*_1C_ = 0) during the mixing time, only the ^13^C’ signal remains and other carbon signals become invisible since no magnetization transfer takes place during the mixing time. On the other hand, when a spin-lock field was applied to the ^13^C channel (e.g., *ν*_1C_ = 76 kHz), the C_*α*_ signal at 54.2 ppm appeared, indicating the magnetization transfer from ^13^C’ to ^13^C_*α*_ during the mixing time. Unlike the static case, where the magnetization can be effectively transferred over the entire molecule through the cross relaxation driven spin diffusion [20], the magnetization transfer under MAS appears to be highly selective, only to its bonded/nearest C_*α*_ nuclei, when the spin-lock field is applied only on the ^13^C channel.

**Fig. 2.**
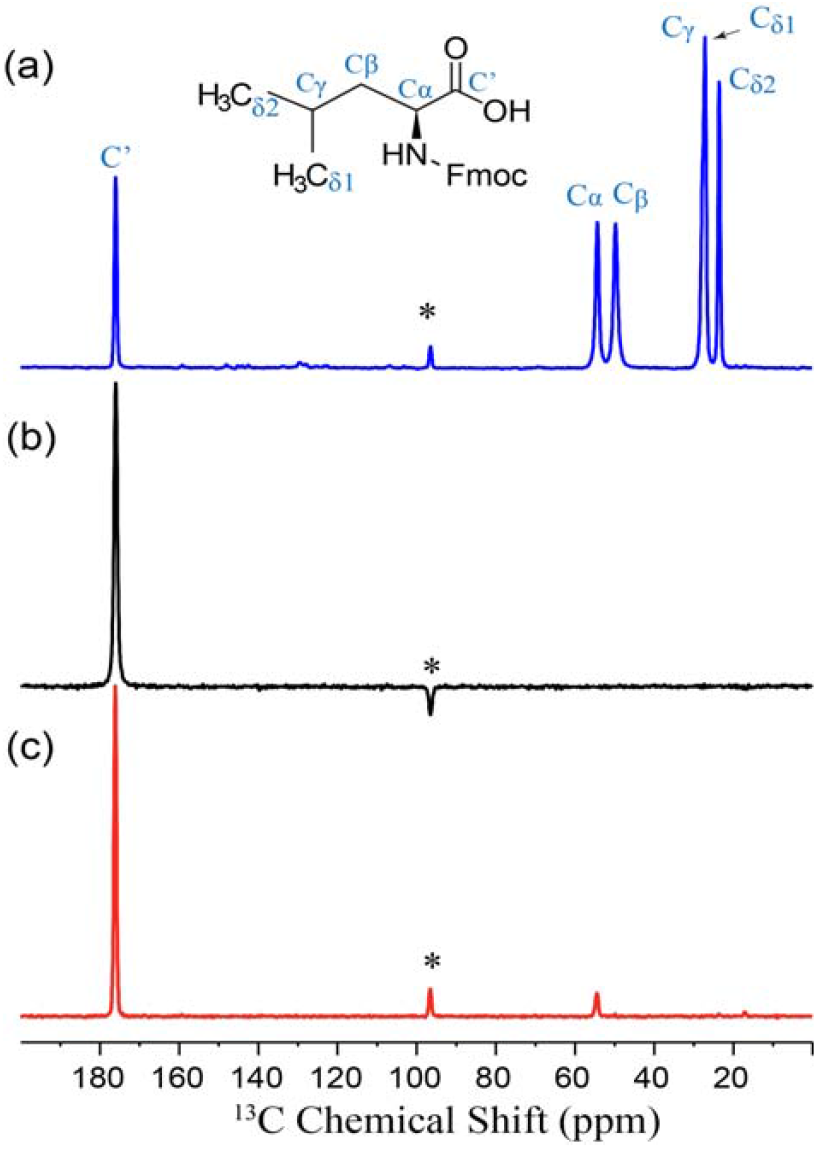
^13^C NMR spectra of a powder sample of Fmoc-leucine obtained under 16 kHz MAS. (a) CPMAS spectrum showing all ^13^C resonances. (b) and (c) are spectra obtained using the pulse sequence shown in Fig. 1a with (b) *ν*_1C_ = 0 and (c) 76 kHz, while with *ν*_1H_ = 0 and. A mixing time, t_mixing_, of 2 ms was used. The asterisks indicate the spinning sideband from ^13^C’ resonance.

Next, ^13^C_*α*_ signal intensity at 54.4 ppm was measured as a function of the ^13^C spin-lock amplitude (ν_1C_) for a given ^1^H spin-lock amplitude (ν_1H_). As indicated in Fig. 3, the observed ^13^C_*α*_ signal intensity gradually increases as *ν*_1C_ increases, when *ν*_1H_ was set to 0 and 16 kHz, under the rotary resonance condition [31]. In order to avoid using the spin-locking under the rotary resonance conditions [32], we set *ν*_1H_ to half integers of the spinning rate, i.e., 1.5ν_r_ (24 kHz), 2.5ν_r_ (40 kHz), 3.5ν_r_ (56 kHz), and 4.5ν_r_ (72 kHz). As shown in the profiles in Fig. 3, it appears that the maximum ^13^C_*α*_ signal intensity was observed at around (ν_1C_-ν_1H_) = ±ν_r_, where *ν*_r_ is the sample spinning rate. For instance, when *ν*_1H_=3.5ν_r_ (56 kHz), the maximum ^13^C_*α*_ signal intensity appears at (ν_1C_-ν_1H_) = 18 kHz, i.e. *ν*_1C_=74 kHz (∼ 4.5ν_r_). While at *ν*_1H_=4.5ν_r_ (72 kHz), the maximum transferred ^13^C_*α*_ signal intensity takes place at (ν_1C_-ν_1H_) = -14 kHz, i.e. *ν*_1C_=58 kHz (∼ 3.5ν_r_). The maximum ^13^C_*α*_ signal intensity obtained under these conditions is almost twice the signal obtained at *ν*_1C_=80 kHz when *ν*_1H_=0 or 16 kHz. It is also clear that the profiles shown in Fig. 3 are similar to the typical cross polarization Hartmann-Hahn matching condition that is broken into a series of sidebands at Δ=±nν_r_ under MAS, where Δ is the mismatch between the two spin-lock RF fields and n is an integer number [33-34]. It can be noticed from Fig. 3 that a stronger *ν*_1C_ gives rise to a more transferred ^13^C_*α*_ signal intensity.

**Fig. 3.**
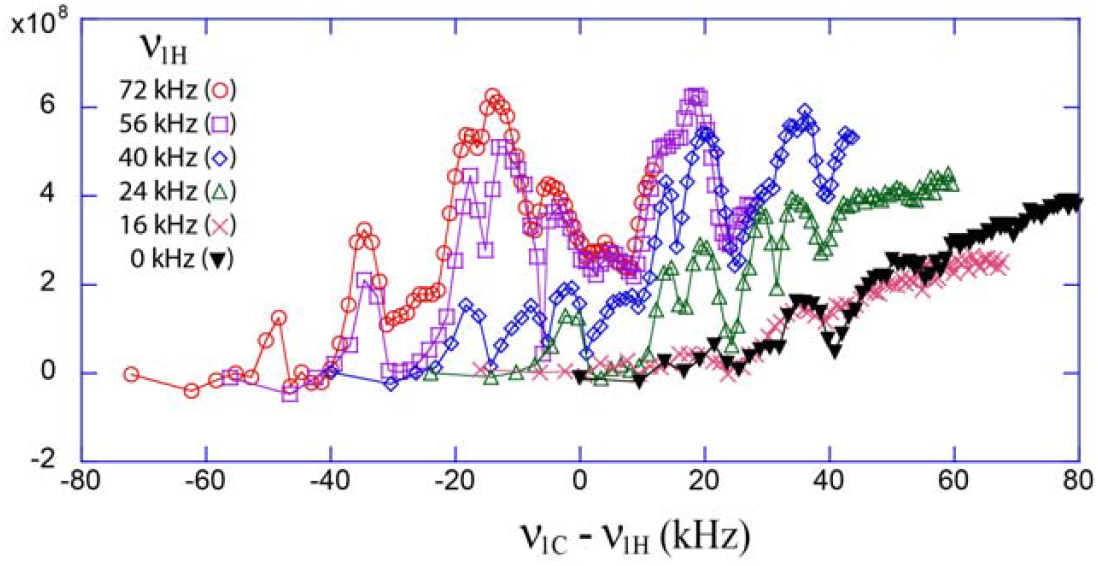
Experimentally measured ^13^C_*α*_ signal intensity transferred from ^13^C’ in Fmoc-L-leucine using the pulse sequence shown in Fig. 1a as a function of (ν_1C_ - *ν*_1H_) at a given *ν*_1H_ as listed in the figure, where *ν*_1C_ was varied from 0 to 80 kHz. The mixing time, t_mixing_, was set to 2 ms.

In order to investigate the magnetization transfer dynamics from ^13^C’ to other carbons in the Fmoc-L-leucine sample, we examined the ^13^C signals intensities as a function of t_mixing_ under the following optimal conditions: i) *ν*_1C_=80 kHz and *ν*_1H_=0; ii) *ν*_1C_ ∼ 3.5ν_r_ (58 kHz) and *ν*_1H_ ∼ 2.5ν_r_ (40 kHz), and iii) *ν*_1C_ ∼ 3.5ν_r_ (58 kHz) and *ν*_1H_ ∼ 4.5ν_r_ (72 kHz). When *ν*_1C_=80 kHz and *ν*_1H_=0, the ^13^C_*α*_ signal intensity built up steadily as a function of t_mixing_, while no other signals were observed, as shown in Fig. 4a. Clearly, under MAS condition, the magnetization transfer is highly selective and appears to take place only between the directly bonded carbons, when there is no spin-lock field on the ^1^H RF channel. This observation is very different from that reported for the static case where the magnetization transfer can be very effective over the entire molecule through the cross relaxation driven spin diffusion [20]. On the other hand, the magnetization transfer under MAS was found to be highly selective, only to its bonded ^13^C_*α*_ spin, when the spin-lock field is applied only on the ^13^C RF channel.

**Fig. 4.**
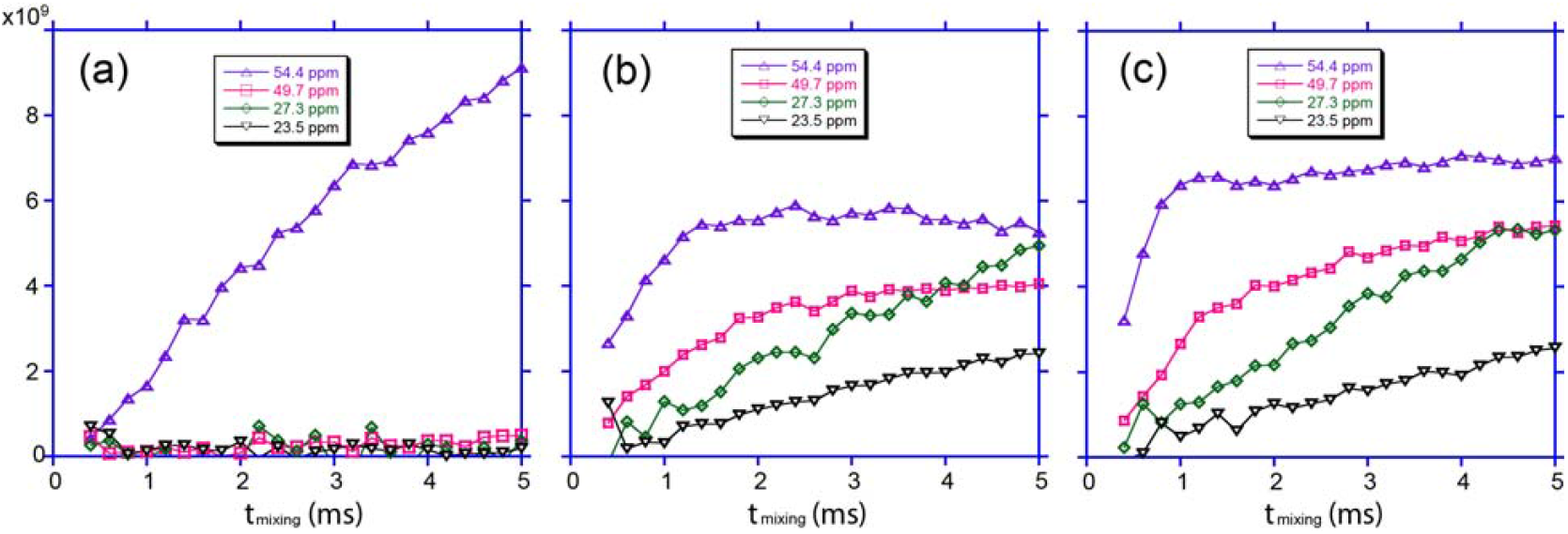
Experimentally observed ^13^C signal intensities from a Fmoc-L-leucine powder sample obtained using the pulse sequence shown using Fig. 1a as a function of t_mixing_ under different conditions. (a) *ν*_1C_=80 kHz and *ν*_1H_=0; (b) *ν*_1C_ ∼ 3.5ν_r_ (58 kHz) and *ν*_1H_ ∼ 2.5ν_r_ (40 kHz); and (c) *ν*_1C_ ∼ 3.5ν_r_ (58 kHz) and *ν*_1H_ ∼ 4.5ν_r_ (72 kHz). The chemical shifts at 54.4, 49.7, 27.3, and 23.5 ppm correspond to C_*α*_, C_β_, C_*γ*_/C_δ1_, and C_δ2_, respectively.

However, when *ν*_1C_ ∼ 3.5ν_r_ (58 kHz) and *ν*_1H_ ∼ 2.5ν_r_ (40 kHz), i.e. (ν_1C_ - *ν*_1H_) = +ν_r_, the ^13^C’ magnetization effectively transfers not only to ^13^C_*α*_, but also to other carbons as well. It can be noticed from Fig. 4b that the ^13^C_*α*_ signal built up rapidly, reaching a plateau at about 1.2 ms and then decayed very slowly. In contrast, the ^13^C_β_ signal built up slower than that of ^13^C_*α*_, but faster than that of ^13^C_δ2_. These observations suggest that the magnetization transfer rate is distance dependent. For the signals at 27.3 ppm containing ^13^C_*γ*_/^13^C_δ1_, their buildup rates were found to be between that observed for ^13^C_β_ and ^13^C_δ2_, behaving just like a simple sum of the buildup curves for ^13^C_β_ and ^13^C_δ2_.

For comparison, we chose the same *ν*_1C_ ∼ 3.5ν_r_ (58 kHz), but used a higher *ν*_1H_ ∼ 4.5ν_r_ (72 kHz), i.e. (ν_1C_ - *ν*_1H_) = -ν_r_. Under this condition, the magnetization transfer rates from ^13^C’ to other carbons, except ^13^C_δ2_, were boosted further, as shown in Fig. 4c. The observed ^13^C_*α*_ signal reached a plateau at about 1.0 ms. Fig.5 shows the plot of stacked 1D spectra and the percentages of all resonances as a function of mixing time. Since the total signal intensity is subject to decay under the spin-lock field, a standard relaxation fitting yielded *T*_1ρ_ = 82.35 ± 7.58 ms, while the ^13^C’ signal at 176.0 ppm decayed at a rate of 12.03 ± 0.62 ms. This large difference in the observed *T*_1ρ_ values is the result of the magnetization transfer from ^13^C’ to other carbons during the mixing time, which can be characterized approximately by a simple model [35]:

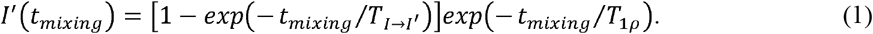

In this equation, T_1*ρ*_ represents the relaxation time under the spin-lock field, and 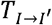 characterizes the magnetization transfer time from one spin to another, in our case, from ^13^C’ to other carbons. Using *T*_1ρ_ = 82.35 ± 7.58 ms as a constant, we fitted the ^13^C buildup curves shown in the bottom right of Fig. 5, yielding 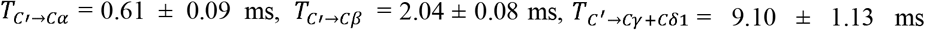, and 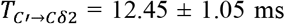. Clearly the magnetization transfer time becomes longer when the carbon position is farther away from the C’ site, suggesting that it is related to the internuclear distance. In addition, it may also be affected by the mismatch between the two spin-lock RF fields, similar to the standard cross-polarization [36].

**Fig. 5.**
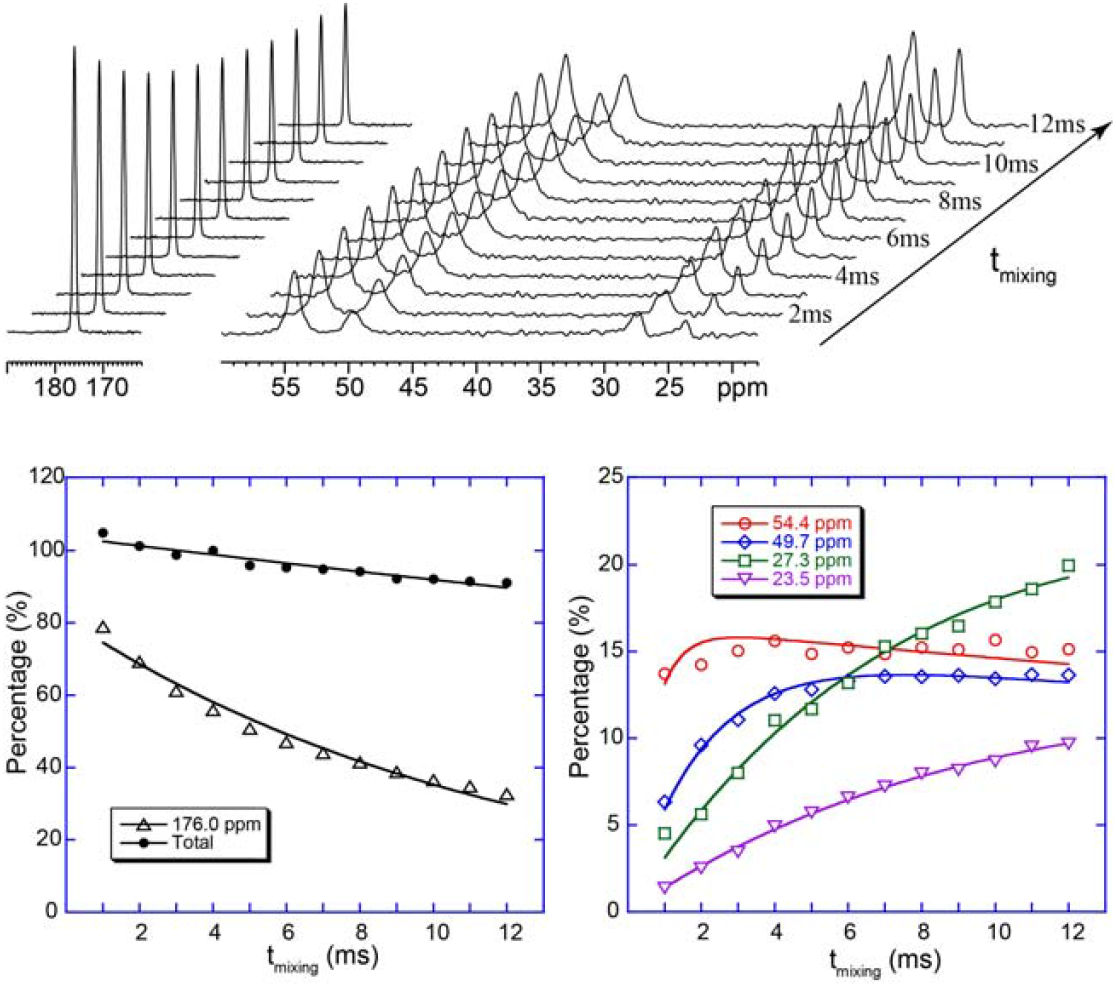
(Top) Plot of stacked 1D NMR spectra of a Fmoc-leucine powder sample obtained at different t_mixing_ under the condition of *ν*_1C_ ∼ 3.5ν_r_ (58 kHz) and *ν*_1H_ ∼ 4.5ν_r_ (72 kHz). (Bottom) Relative signal intensities as a function of t_mixing_. The total intensities were normalized to that obtained for t_mixing_=4 ms. The data in the bottom left were fitted through a standard *T*_1ρ_ procedure, while the data in the bottom right were fitted using Eq.(1) with the *T*_1ρ_ value of 82.35 ms as a constant obtained from the total signal fitting in the bottom left.

Based on the experimental optimization above, we conclude that a remarkably rapid ^13^C spin-exchange process takes place under MAS when this mismatch condition (ν_1C_ - *ν*_1H_) = ±ν_r_ is fulfilled. Next, we applied this mismatch condition to ^15^N nuclei in order to examine whether it provides a sufficient ^15^N spin-exchange where ^15^N-^15^N dipolar coupling is extremely weak. Fig. 6 shows the 2D ^15^N-^15^N homonuclear chemical shift correlation spectra of a powder sample of ^15^N-labeled L-histidine amino acid (pH 6.3) using the RF pulse sequence shown in Fig. 1b. For comparison, the standard homonuclear correlation experiment using dipolar assisted rotational resonance (DARR) during the mixing time [31, 37] was also performed. A total of six ^15^N resonances at 250.0, 190.6, 177.4, 171.9, 48.6 and 42.2 ppm were observed due to the coexistence of charged- and *τ*-tautomers at pH 6.3.[38-39] It is not surprising that, as shown in Fig. 6a, ^15^N-^15^N DARR spectrum does not yield any cross peaks even with a mixing time of 50 ms due to the extremely weak ^15^N-^15^N dipolar couplings within the two histidine tautomers. However, when using the RF pulse sequence shown in Fig. 1b under the mismatch condition (ν_1N_ - *ν*_1H_) = -ν_r_, two sets of cross peaks were clearly observed even with a mixing time as short as 5.0 ms, in spite of the extremely weak ^15^N-^15^N dipolar couplings. One set of cross peaks are among the resonances of 190.6, 177.4, and 48.6 ppm, Fig. 7 shows the 2D ^15^N-^15^N homonuclear chemical shift correlation spectra of the uniformly-^15^N,^13^C-labeled AqpZ protein in synthetic DMPC lipid bilayers using different experimental schemes. Again, the 2D spectrum obtained with DARR mixing in Fig. 7a shows strong uncorrelated diagonal peaks but no cross peaks at all for a mixing time of 50 ms. However, when using the RF pulse sequence shown in Fig. 1b under the mismatch condition (ν_1N_-ν_1H_) = -ν_r_ for a 5.0 ms mixing time, ^15^N-^15^N correlations were observed throughout the amide-^15^N chemical shift range, as shown in Fig. 7b. It is illustrated by the two 1D spectral slices extracted at 101.4 and 98.9 ppm indicated by the green dashed lines in the 2D spectrum (Fig. 7b) that each Gly ^15^N resonance correlates with two resonances, suggesting that the ^15^N spin exchange primarily takes place between two neighboring residues, i.e., N_i_-N_i±1_, within 5.0 ms mixing time under the mismatch condition (ν_1N_-ν_1H_) = -ν_r_. Unlike the indirect ^15^N correlation scheme through protons, i.e., the pathway of ^15^N → ^1^H → ^1^H → ^15^N, where, after t_1_ evolution, ^15^N magnetization is sent back to ^1^H for spin exchange and brought back for detection in the t_2_ dimension [15, 40-43], the long-range ^15^N correlations occur because the proton driven spin exchange takes place in the proton reservoir from one proton to other protons.

**Fig. 6.**
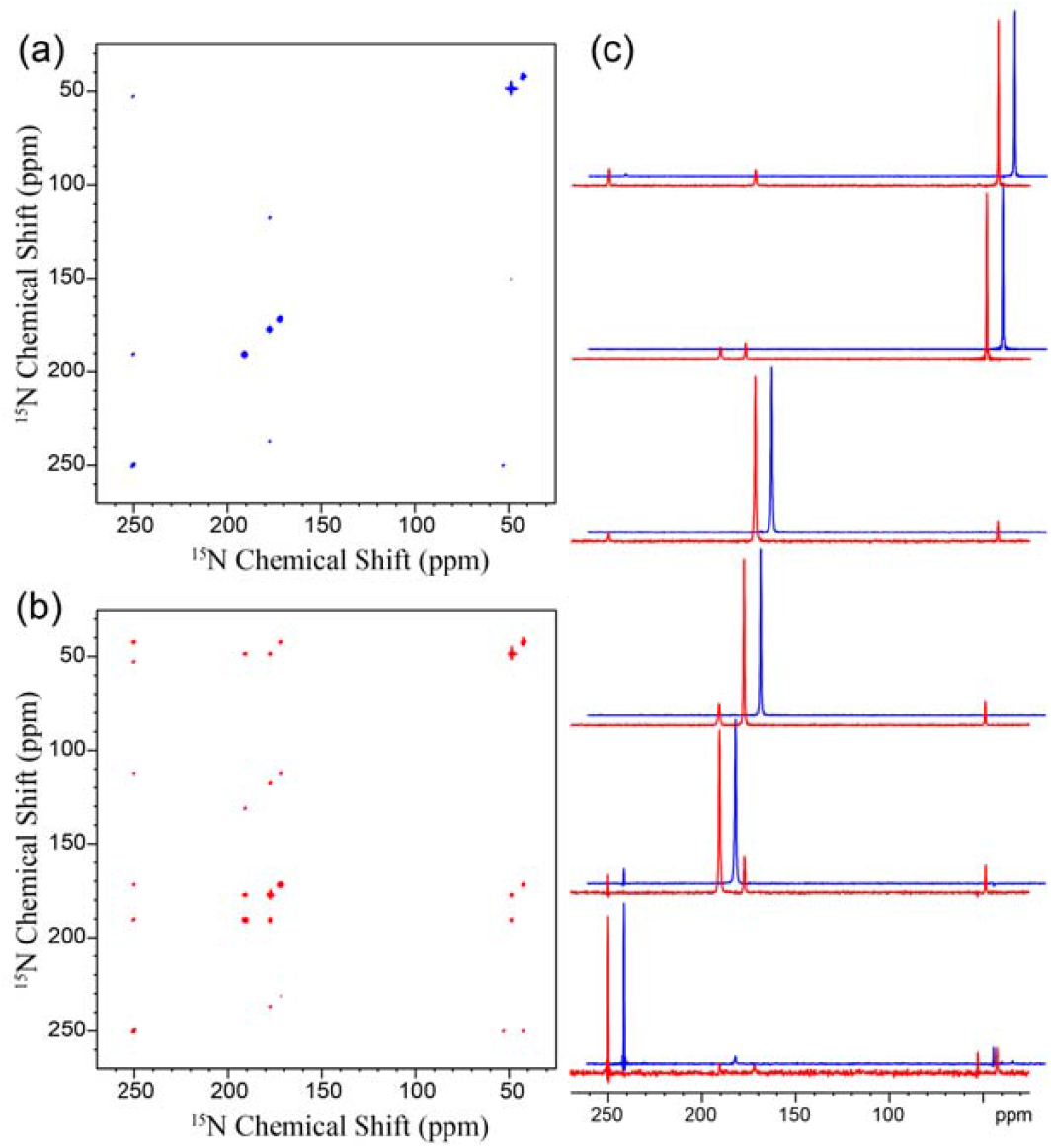
2D ^15^N-^15^N chemical shift correlation spectra of ^15^N-labeled L-histidine amino acid (pH 6.3) powder sample obtained using: (a) the standard correlation experiment with a dipolar assisted rotational resonance (DARR) mixing time of 50 ms, where ^1^H spin-lock field of 16 kHz, i.e., *ν*_r_, was used to enhance ^15^N spin diffusion; (b) the RF pulse sequence shown in Fig. 1b with a mixing time of 5.0 ms where *ν*_1N_ = 3.5ν_r_ (58 kHz) and *ν*_1H_ = 4.5ν_r_ (72 kHz). (c) comparison of 1D spectral slices taken at 250.0, 190.6, 177.4, 171.9, 48.6, and 42.2 ppm (from bottom to which belongs to the charged tautomer, while the other one among the peaks between 250.0, 171.9, and 42.2 ppm, representing the *τ*-tautomer. The 1D spectral slices taken along each of the resonances at 250.0, 190.6, 177.4, 171.9, 48.6, and 42.2 ppm clearly show the presence of cross peaks from Fig. 6b (red) and the absence of cross peaks from Fig. 6a (blue). Such an ability to directly correlate ^15^N resonances within each tautomer opens up an efficient avenue to identify histidine tautomeric states at different pH in complex biomolecular systems.

**Fig. 7.**
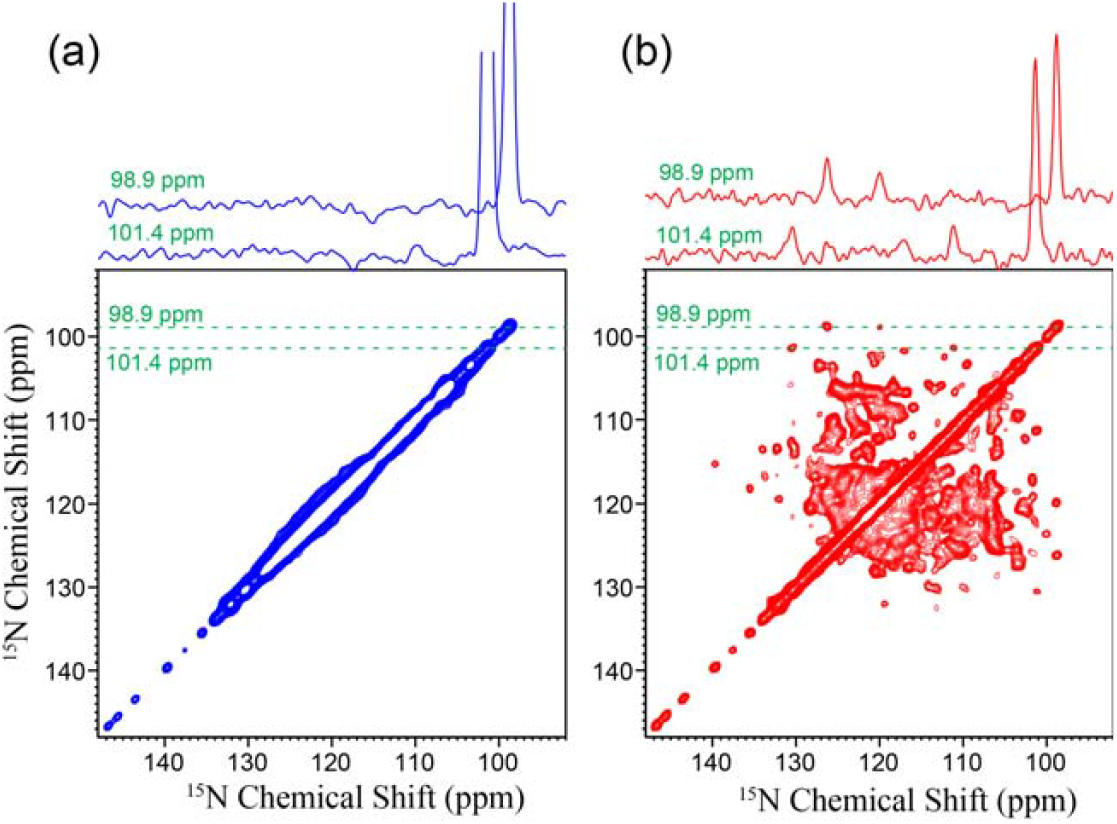
2D ^15^N-^15^N chemical shift correlation spectra of a uniformly ^15^N,^13^C labeled AqpZ protein reconstituted in DMPC lipid bilayers using (a) the standard correlation experiment with a dipolar assisted rotational resonance (DARR) mixing time of 50 ms, where ^1^H spin-lock field of 16 kHz, i.e., *ν*_r_, was used to enhance ^15^N spin diffusion; (b) the pulse sequence shown in Fig. 1b, with a mixing time of 5.0 ms during which *ν*_1N_ = 3.5ν_r_ (58 kHz) and *ν*_1H_ = 4.5ν_r_ (72 kHz). On the top of the spectra are the slices extracted along the dashed lines (98.9 and 101.4 ppm). In the experiments, 64 scans were used to accumulate the signals for each t increment.

In the resonance-assignment strategies for biological systems, the N-C heteronuclear correlations are crucial for sequential assignments due to the fact that each backbone nitrogen is covalently bonded to the carbonyl carbon of i^th^ residue and the C_α_ of (i+1)^th^ residue providing a through-bond linkage between the two sequential residues. Thus, three dimensional (3D) schemes such as the so-called CANCO experiments [44-46] are used to establish the connectivity between the two neighboring residues through the correlation between their respective ^13^C’ and ^13^C_α_ resonances. The ability to correlate the ^15^N resonances between two neighboring residues, as demonstrated here, opens up an attractive way to establish the connectivity between neighboring residues in a polypeptide or protein, especially when combining with ^1^H-detection, for resonance assignments.

## Conclusion

Here, we demonstrate that applying double spin-lock RF pulses enables efficient magnetization exchange among low-γ nuclei, thereby allowing the acquisition of 2D homonuclear chemical-shift correlation spectra under MAS conditions. In this scheme, the observed magnetization transfer arises from a combination of RF and cross-relaxation assisted spin diffusion processes, both of which contribute to overcoming the inherently weak dipolar couplings among low-γ nuclear spins such as ^13^C and ^1^□N. To better understand the underlying mechanisms, we experimentally characterized how the magnetization-exchange efficiency depends on two key parameters: the MAS frequency, which modulates the dipolar interactions, and the Hartmann– Hahn mismatch, which governs the effectiveness of the spin-lock fields in mediating the magnetization exchange. These measurements confirm that both factors play significant roles in shaping the transfer dynamics, providing insight into how RF-driven cross-relaxation and MAS-modulated dipolar couplings work together to produce the observed chemical shift correlations. Perhaps the most striking demonstration of the power of this sequence comes from experiments on uniformly ^1^LN-labeled aquaporin reconstituted in DMPC lipid vesicles. Under standard DARR mixing, a 50 ms period fails to generate any detectable 2D ^15^N–^15^N cross peaks, reflecting the intrinsic inefficiency of conventional dipolar-driven exchange for ^15^N nuclei due to the very weak ^15^N–^15^N dipolar couplings. In contrast, applying a 5.0 ms mixing time with the double spin-lock sequence under the optimized Hartmann–Hahn mismatch condition yields a fully correlated 2D ^15^N–^15^N spectrum, enabling observation of cross peaks that are otherwise unobservable. Remarkably, the ability to selectively correlate the ^15^N resonances between two neighboring residues would be valuable for accomplishing resonance assignments in the structural studies of a protein. This dramatic improvement highlights the potential of the double spin-lock approach as a robust and general method for homonuclear chemical-shift correlation involving low-γ nuclei in complex biological solids.

## Acknowledgements

All NMR experiments were carried out at the National High Magnetic Field Lab (NHMFL) supported by the NSF Cooperative Agreement DMR-2128556 and the State of Florida. We thank Dr. Huayong Xie and Prof. Jun Yang from Wuhan Institute of Physics and Mathematics/Innovation Academy for Precision Measurement Science and Technology, Chinese Academy of Sciences, for providing the AqpZ protein sample. A.R. acknowledges the funding support from NIH (R35GM13973) and FSU.

## Supplemental Materials

2D ^13^C-^13^C chemical shift correlation spectra of a uniformly ^13^C, ^15^N-labeled Fmoc-L-leucine powder sample under different experimental conditions; Additional 1D spectral slices extracted at 128.2, 125.6, 120.4, and 117.2 ppm from the 2D ^15^N-^15^N chemical shift correlation spectrum in Fig. 7.

