## Supporting Information for "Homonuclear Chemical Shift Correlation in Solids Under MAS by Fast Cross-Relaxation Driven Spin Diffusion"

Figure 1S shows 2D ^13^C-^13^C chemical shift correlation spectra of a uniformly ^13^C, ^15^N-labeled Fmoc-L-leucine powder sample under different experimental conditions. With a 5 ms DARR mixing time, the 2D spectrum (Top left) shows the ^13^C correlations between the covalently bonded carbons only, as expected. Similarly, when using the RF pulse sequence shown in Fig. 1b, with the spin-lock field applied only on the ^13^C channel (e.g. ν_1C_ = 80 kHz and ν_1H_ = 0 kHz), the ^13^C/^13^C correlations appear only between the covalently bonded carbons as shown in Fig. 1S (Top right). This observation confirms that the magnetization transfer under MAS is highly selective, only between two bonded ^13^C atoms when the spin-lock field is applied only on the ^13^C RF channel, unlike the static case where the magnetization transfer can be very effective over the entire molecule through the cross relaxation driven spin diffusion process. However, when the Hartmann-Hahn mismatch condition between the two spin-lock RF fields is fulfilled (ν_1C_ - ν_1H_) = ±ν_r_, the magnetization transfer becomes very effective over the entire molecule as illustrated in the spectra shown in Fig. 1S (bottom).

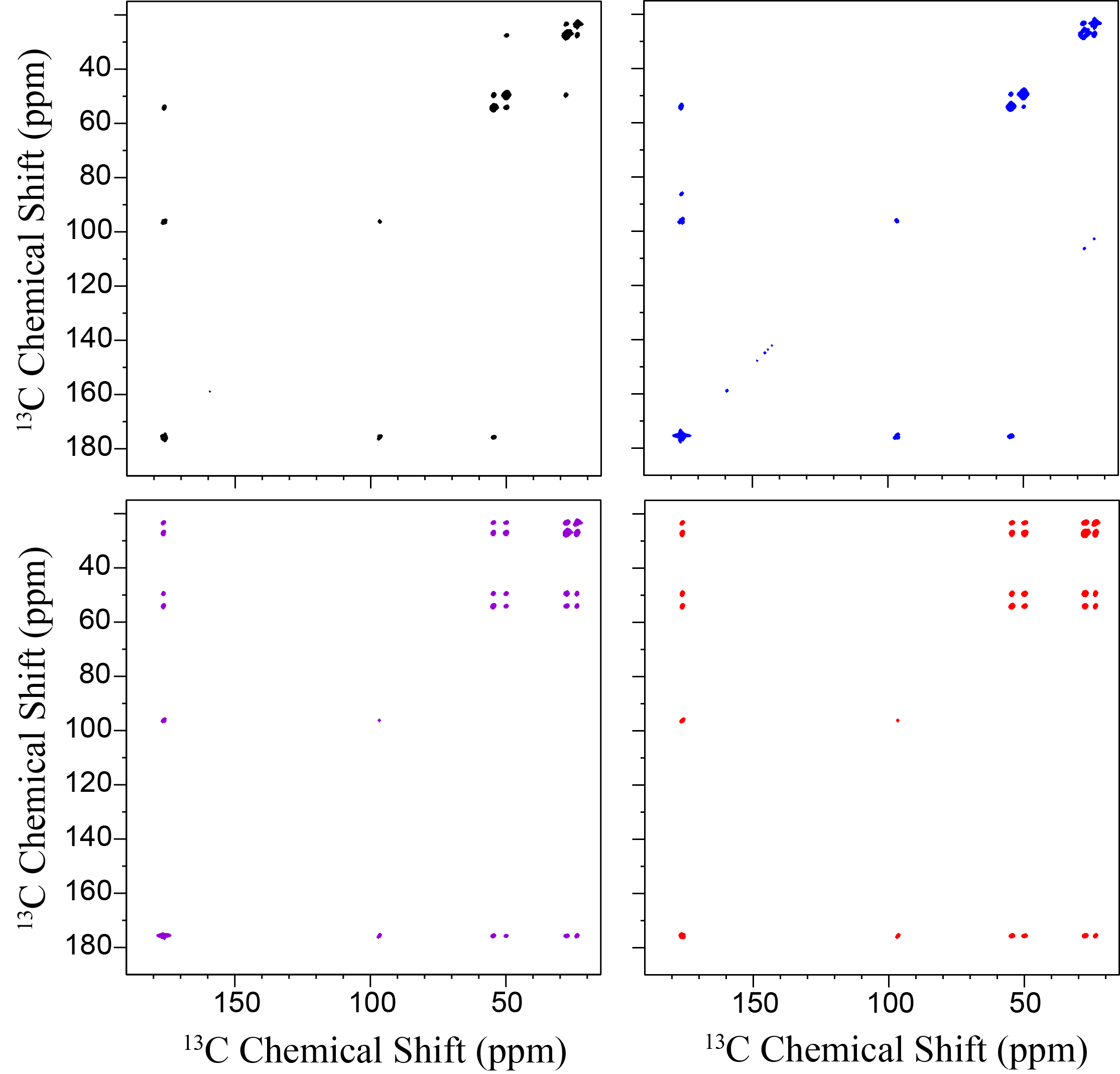

**Figure 1S**. 2D ^13^C-^13^C chemical shift correlation spectra of uniformly ^13^C, ^15^N-labeled Fmoc-L-leucine powder sample obtained using the standard correlation experiment with a dipolar assisted rotational resonance (DARR) mixing where ^1^H spin-lock field of 16 kHz (top left) and the RF pulse sequence shown in Fig. 1b with the RF spin-lock fields of (top right) ν_1C_ = 80 kHz and ν_1H_ = 0 kHz; (bottom left) ν_1C_ ~ 3.5ν_r_ (58 kHz) and ν_1H_ ~ 2.5ν_r_ (40 kHz); (bottom right) ν_1C_ = 4.5ν_r_ (76 kHz) and ν_1H_ ~ 3.5ν_r_ (56 kHz). In these experiments, the mixing time was set to 5.0 ms.

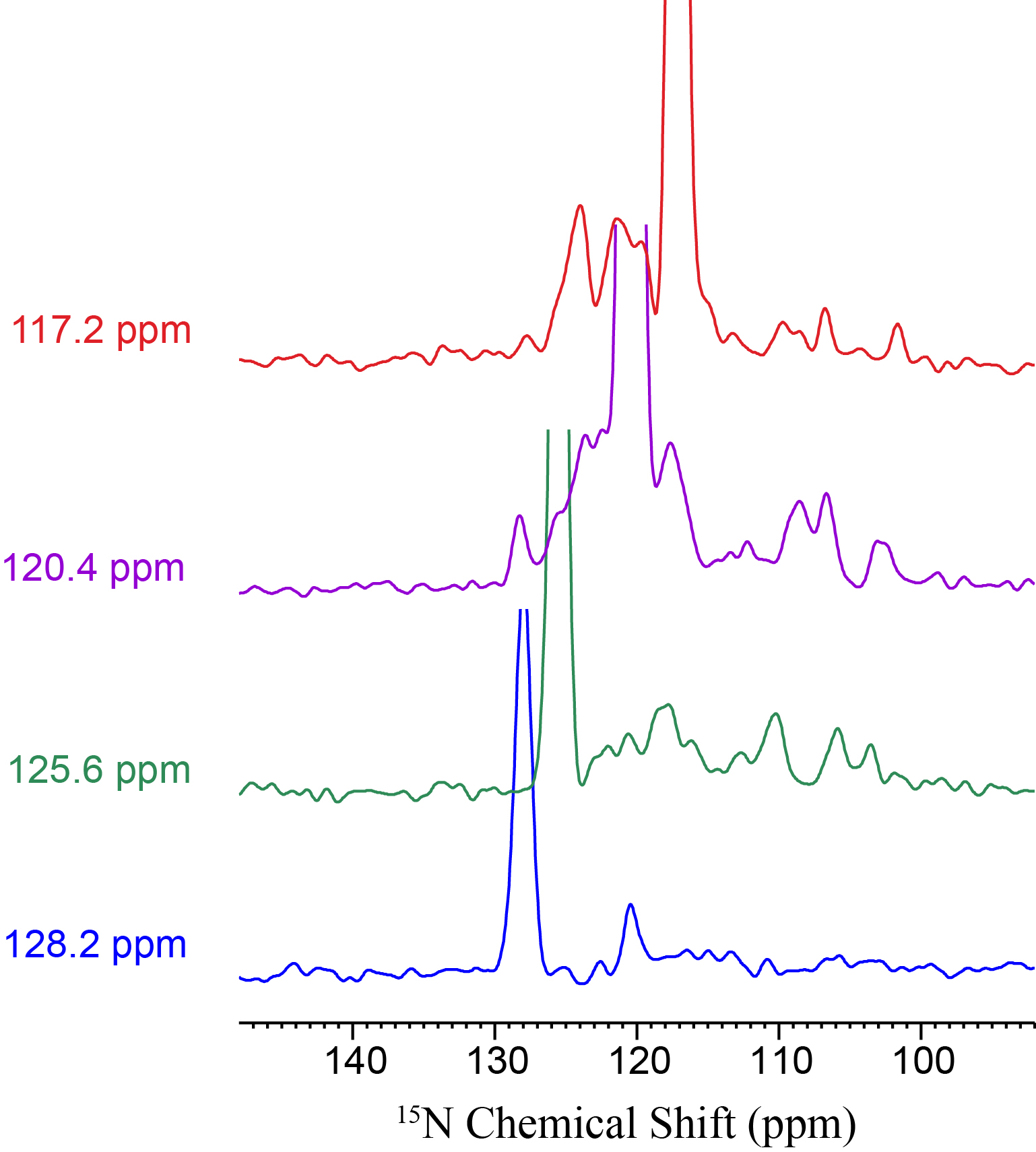

**Fig. 2S** Additional 1D spectral slices exctracted at 128.2, 125.6, 120.4, and 117.2 ppm from the 2D ^15^N-^15^N chemical shift correlation spectrum shown in Fig. 7. The signal intensities are normalized.
